# A model of ganglion axon pathways accounts for percepts elicited by retinal implants

**DOI:** 10.1101/453035

**Authors:** Michael Beyeler, Devyani Nanduri, James D. Weiland, Ariel Rokem, Geoffrey M. Boynton, Ione Fine

## Abstract

Retinal prostheses, now implanted in over 250 patients worldwide, electrically stimulate surviving cells in order to evoke neuronal responses that are interpreted by the brain as visual percepts (‘phosphenes’). However, instead of seeing focal spots of light, current implant users perceive highly distorted phosphenes that vary in shape both across subjects and electrodes. We characterized these distortions by asking users of the Argus retinal prosthesis system (Second Sight Medical Products) to draw elicited percepts on a touchscreen. We found that phosphene shape could be accurately predicted by simulating the topographic organization of nerve fiber bundles in each subject’s retina. Our model shows that activation of ganglion axons contributes to a rich repertoire of phosphene shapes, successfully replicating percepts ranging from ‘blobs’ to oriented ‘streaks’ and ‘wedges’ depending on electrode location. This work provides a first step towards future devices that incorporate stimulation strategies tailored to each individual patient’s retinal neurophysiology.

**One Sentence Summary:** We show that the perceptual experience of retinal implant users can be accurately predicted using a computational model that simulates the topographic organization of each individual patient’s retinal ganglion axon pathways.

## 1. Introduction

Degenerative retinal diseases such as retinitis pigmentosa (*2*) and macular degeneration (*3*) lead to a loss of photoreceptor cells and subsequent remodeling of the neural circuitry in the retina (*4, 5*), causing irreversible blindness in more than 10 million people worldwide. Analogous to cochlear implants, the goal of retinal prostheses is to help alleviate these incurable conditions by electrically stimulating surviving cells in the retina (for a recent review, see 6). The hope is that electrically evoked neuronal responses will be transmitted to the brain and interpreted by the subject as visual percepts (‘phosphenes’).

Two devices are already approved for commercial use: Argus II (epiretinal, Second Sight Medical Products Inc., 7, 8) and Alpha-IMS (subretinal, Retina Implant AG, 9), which have been shown to restore vision up to a visual acuity of 20/1,260 (*10*) and 20/546 (*9*), respectively. In addition, PRIMA (subretinal, Pixium Vision, 11) has started clinical trials, with others to follow shortly. In combination with stem cell therapy (*12, 13*) and optogenetics (*14*), a range of sight restoration options should be available within a decade (*15*).

However, despite the increasing clinical and commercial use of these devices, the perceptual experience of retinal implant users remains poorly understood. For example, even in response to single-electrode stimulation, the appearance of individual phosphenes is highly variable not only across subjects but also across electrodes within a subject, with subjects typically reporting seeing distorted and often elongated geometric shapes that fade quickly over time (*16–23*). Furthermore, linearly combining these ‘building blocks’ of percepts from individual electrodes often fails to predict the combination of percepts evoked when multiple electrodes are stimulated (*18, 24–26*). Consequently, most subjects cannot determine the orientation of gratings that are used to measure visual acuity, and those who can recognize letters take more than 40 seconds to do so (*27, 28*).

Both computational (*29, 30*) and *in vitro* electrophysiological studies (*Fried et al., 2009; Rizzo et al., 2003; Weitz et al., 2015*) suggest that electrode configurations similar to those implanted in patients do not achieve focal activation, but rather produce significant activation of passing axon fibers, which may result in perceptual distortions in patients. Here, we are the first to directly examine whether axonal stimulation contributes to the rich repertoire of phosphene shapes reported by patients. Our computational model can account for the apparent shape of phosphenes elicited by single-electrode stimulation in two generations of the Argus retinal prosthesis system (Second Sight Medical Products, Inc.).

Four subjects suffering from severe retinitis pigmentosa were chronically implanted with an epiretinal prosthesis in the macular region of the retina: one subject was implanted with an Argus I device (16 platinum disc electrodes arranged in a 4×4 checkerboard pattern; see **Figure 1A**), and three subjects were implanted with Argus II device (60 platinum disc electrodes in a 6×10 arrangement; **Figure 1B**). Electrical stimulation was delivered to a number of pre-selected electrodes in random order (five repetitions each) using square-wave, biphasic, cathodic-first pulse trains with fixed stimulus duration, and we asked subjects to outline perceived phosphene shape either on a grid screen (Argus I; **Figure 1B, C**) or a computer touchscreen (Argus II; **Figure 1E, F**) (see Methods). We then used a computational model to generate predictions about the apparent shape of the expected visual percepts, and compared the predicted images to patient phosphene drawings. The model assumed that distortions are due to activation of ganglion axon pathways, having estimated the spatial layout of these pathways using traced nerve fiber bundle trajectories extracted from ophthalmic fundus photographs of 55 human eyes (*1*).

**Figure 1:**
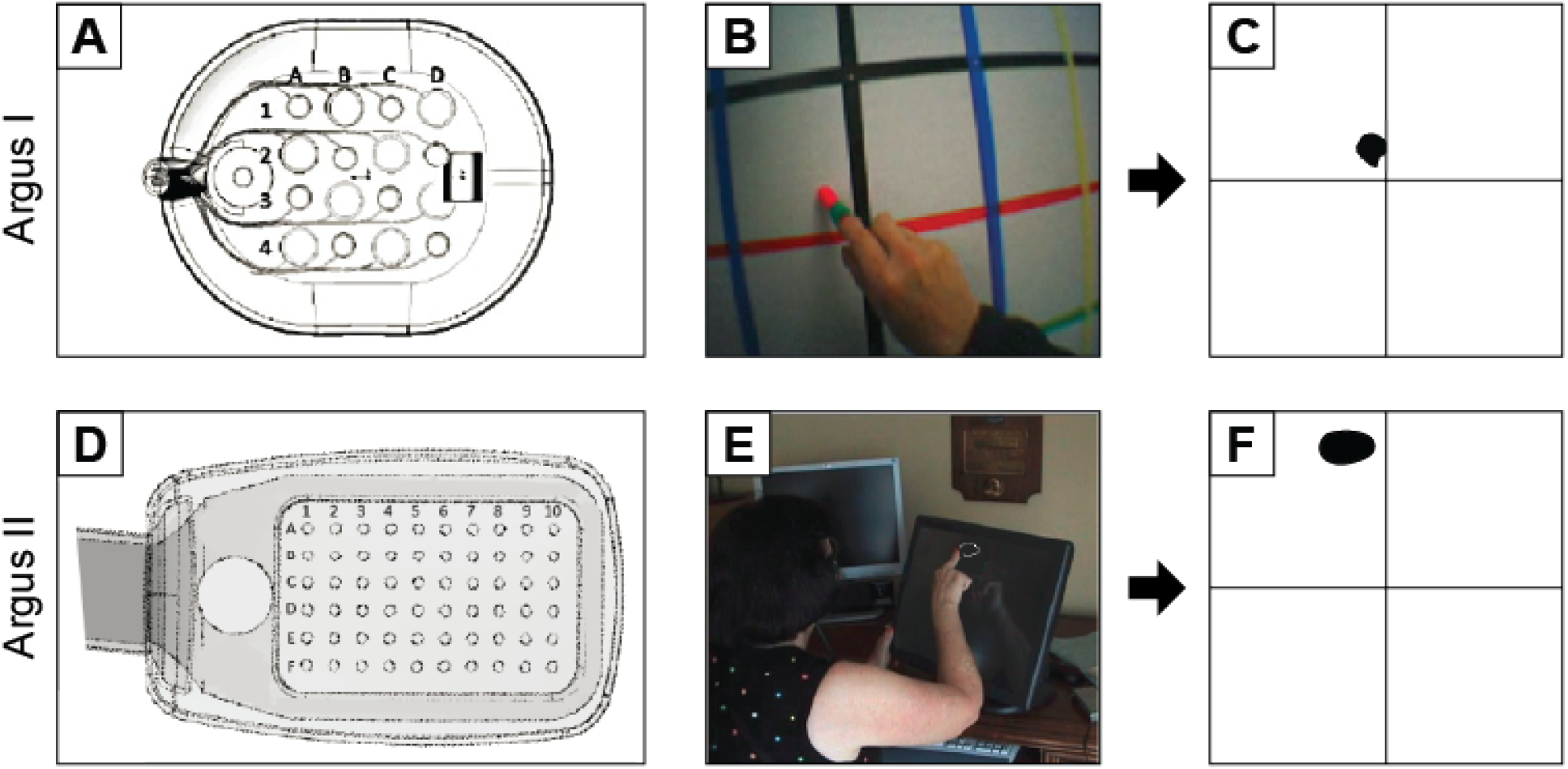
Retinal implants used for the drawing task. (**A**) Argus I electrode array (4×4 electrodes of 260 μm or 520 μm diameter arranged in a checkerboard pattern). (**B**) Argus I subject drawings on a grid screen were captured by an external camera and recorded to a video file. (**C**) Video files were analyzed offline by tracking the location of the fingertip frame-by-frame and by translating the drawings to a binary image. (**D**) Argus II electrode array (6×10 electrodes of 200 μm diameter). (**E**) Argus II subject drawings were recorded by a touch screen monitor. (**F**) Subject drawings were translated to a binary image. Shapes were closed by automatically connecting the first and last tracked fingertip location, after which a floodfill was applied.

## 2. Results

### 2.1 Phosphene drawings vary across electrodes, but are relatively consistent for a given electrode

All subjects consistently reported seeing phosphenes upon electrical stimulation of the retina. Phosphenes appeared light gray, white, or yellowish in color. However, phosphene drawings varied greatly across subjects and electrodes; representative drawings for each subject are shown in **Figure 2.** Whereas stimulation of some electrodes elicited consistent percepts across trials (top row of panels in each subplot), stimulation of other electrodes led to percepts that varied in both size and shape across trials (bottom row of panels in each subplot). Subjects occasionally but rarely reported seeing two distinct shapes (e.g., Subject 1, Electrode D1). Mean images for each electrode were obtained by averaging the drawings across the five stimulation trials, aligned by their centers of mass (column ‘average’). Mean images were then centered over the corresponding electrode in a schematic of the subject’s implant to reveal the rich repertoire of elicited percepts across electrodes (large, rightmost panel in each subplot).

**Figure 2:**
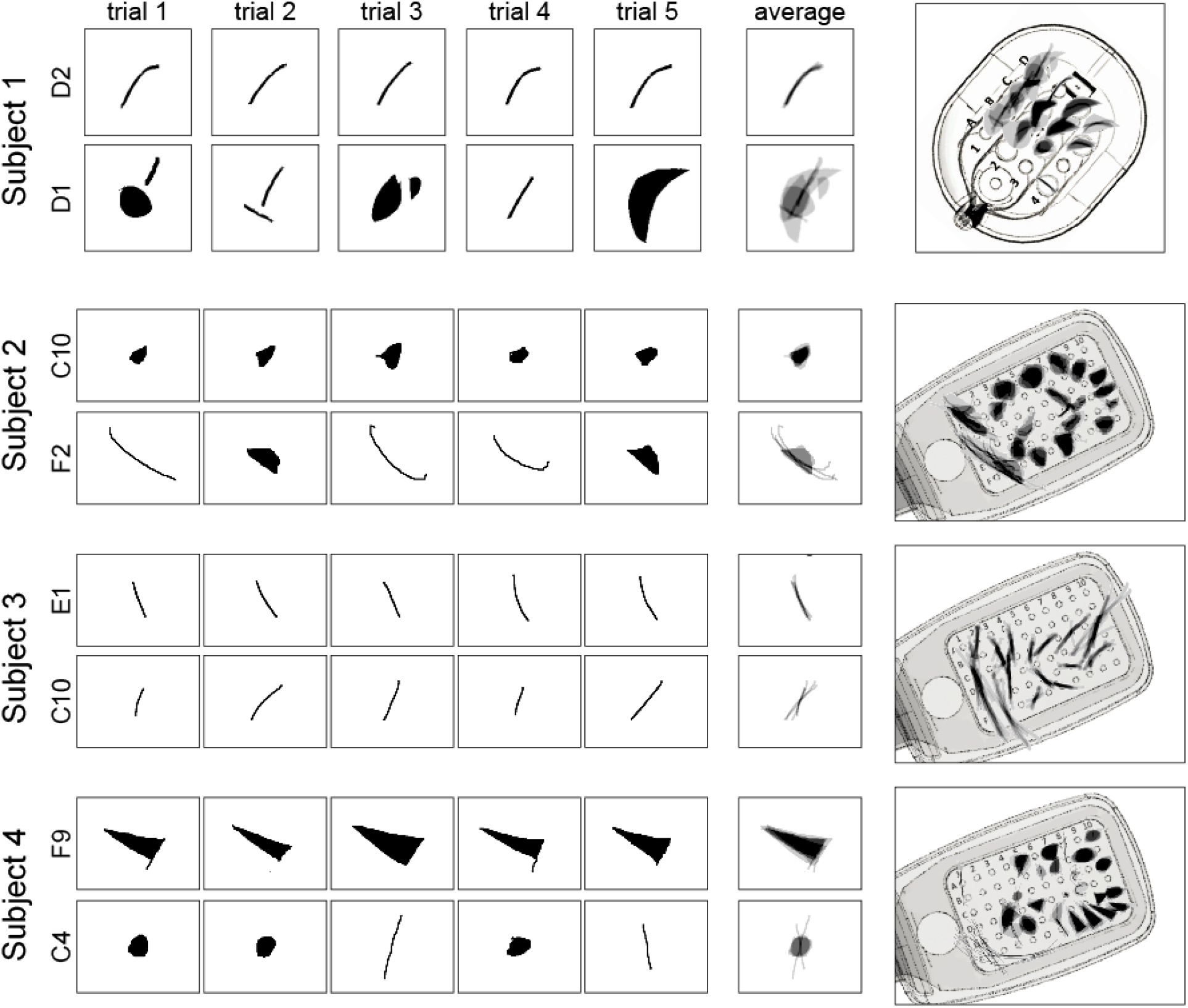
Phosphene drawings vary across electrodes. Drawings from individual trials are shown for the most consistent (top row in each panel) and least consistent electrodes (bottom row in each panel) for Subjects 1–4. Mean images (labeled ‘average’) were obtained by averaging drawings from individual trials aligned at their center of mass. These averaged drawings were then overlaid over the corresponding electrode in a schematic of each subject’s implant (rightmost column).

As is evident from these data, only a small number of phosphenes could be described as focal spots of light. Subject 1 drew percepts as either curved or straight lines, wedges, or relatively round spots. Subject 2 drew most percepts as ovals or relatively round spots with only a few curved or straight lines of varying thickness, whereas Subject 3 drew all phosphenes as slightly curved or straight, thin lines. Subject 4 predominantly drew ovals, wedges, and triangles, with only few curved or straight lines.

Interestingly, for Subjects 2–4 the percepts produced by electrodes in the first two rows of the array (i.e., Electrodes A1–F1, A2–F2) were much thinner and longer than for other electrodes (*31, 32*). It is possible that these electrodes were the ones that were closest to the retinal surface, since the surgical tack used to attach the implant to the retina was located next to the first row of electrodes. However, we did not have access to optical coherence tomography data, which would have allowed us to directly measure electrode-retina distance.

To quantify the similarity and variability of individual phosphene drawings, we calculated three shape descriptors for each collected drawing: phosphene *area, orientation*, and *elongation* (see Methods). These parameter-free metrics were based on a set of statistical quantities known as ‘image moments’; that is, particular weighted averages of pixel intensities across an image (Equation 1). Phosphene orientation and elongation were calculated from the eigenvalues and eigenvectors of each drawing’s covariance matrix (Equations 2 – 4).

The upper panels of **Figure 3** show distributions of phosphene *area* (**Figure 3A**), *orientation* (**Figure 3B**), and *elongation* (**Figure 3C**) for each subject, across all tested electrodes. The lower panel (**Figure 3D–F**) boxplots depicts trial-to-trial variability for each shape descriptor of a given electrode, measured as the standard error of the mean (SEM) calculated across drawings.

**Figure 3:**
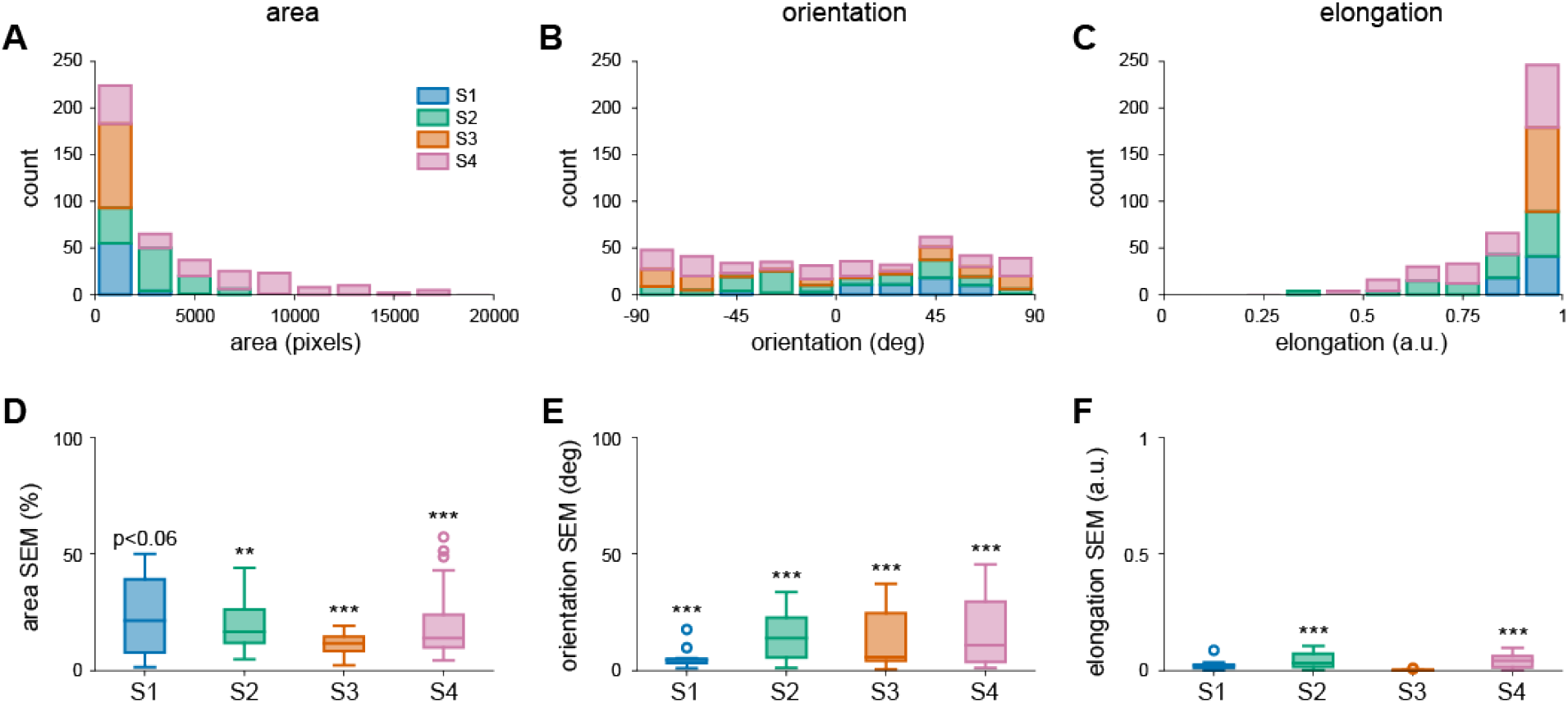
(**A-C**) Distribution of phosphene area, orientation, and elongation for each subject (Subject 1: 60 drawings, Subject 2: 110 drawings, Subject 3: 90 drawings, Subject 4: 140 drawings). (**D-F**) Distribution of the variability of shape descriptors for each subject, measured as the standard error of the mean (SEM) across trials for every electrode. Each box extended from the lower to upper quartile values of the data, with a line at the median. Whiskers extended from the fifth to ninety-fifth percentiles, with data points outside that range considered outliers (‘o’). Area SEM for every electrode was normalized by the mean area of all drawings for that particular electrode.

To assess whether observed SEM values were smaller than would be predicted from a random sample of phosphene drawings we performed a resampling analysis (1000 iterations). We began by calculating the SEM across all five drawings for each electrode. To assess whether, for an individual subject, drawings were more similar for an individual electrode then across other electrodes in that particular subjects’ array, resampling was done by randomly sampling (with replacement) phosphene drawings across all the electrodes of that subject. Probability values were estimated by comparing the mean SEM across electrodes of the real distribution to the 1-tailed confidence interval generated by resampling. Detailed results for each shape descriptor are given below.

### 2.1.1 Phosphene area estimates show consistent variation across electrodes

Phosphene *area* was calculated as the number of nonzero pixels in the drawing. We report data in terms of pixels because the relationship between drawing area and the size of the phosphene in terms of visual angle should be interpreted with caution. Based on each subject’s viewing distance, the touchscreen should have subtended 73.8 × 73.8° of visual angle for Subject 1, 60 × 45° for Subject 2, 65 × 48.8° for Subject 3, and 64 × 48° for Subject 4. Although we asked subjects to draw phosphenes ‘as if they appeared at arm’s length’ (see Methods), subjects qualitatively reported that phosphenes could appear as close as ‘in front of their face’ to ‘at arm’s length’. As can be seen by the variability in the boxplots in **Figure 3D**, estimates of area varied widely across both subjects and electrodes. However, despite the lack of a reference plane in depth, for all but S1 (marginally significant) the observed SEM values for phosphene area were significantly smaller than SEM values from randomly chosen electrodes (S1: *p* < 0.06, S2: *p* < 0.01, S3: *p* < 0.001, S4: *p* < 0.001). S1 had particularly small percepts, so areal variability may have been more heavily influenced by drawing error.

### 2.1.2 Phosphene orientation is more consistent within than across electrodes

Phosphene *orientation* was calculated as the angle of the principal eigenvector (in the range [-90°, 90°]). For all subjects, mean SEM values for phosphene orientation were significantly smaller than mean SEM values for our bootstrapped null model (S1: *p* < 0.001, S2: *p* < 0.001, S3: *p* < 0.001, S4: *p* < 0.001), showing that the variation in phosphene orientation within an individual electrode is less than the variation across electrodes (**Figure 3E**).

### 2.1.3 Phosphene elongation is more consistent within than across electrodes

Phosphene *elongation* was calculated as the relative difference in magnitude of the eigenvalues and normalized to [0, 1], with 0 representing a circle, and 1 representing an infinitesimally thin line. The distribution of elongation values indicates that subjects consistently saw elongated percepts instead of focal spots of light, indeed Subject 3 reported exclusively seeing thin curved and straight lines. These results are in stark contrast to the prevailing assumption in the field that stimulating a single electrode should generate the percept of a small focal spot of light (*33–37*).

For Subjects S2 and S4, observed SEM values for phosphene elongation were significantly smaller than of our bootstrapped null model (S2: *p* < 0.001, S4: *p* < 0.001). For Subjects S1 and S3 results were not significant; percepts were heavily elongated for every electrode, providing little variability in the dataset (**Figure 3F**).

### 2.1.4 Drawing accuracy

Our subjects had been lacking tactile-visual feedback for many years. This motivated a control experiment, where we asked subjects to explore various tactile targets (made of felt with a cardboard background) with their hands, and to draw the targets on a touch screen (**Figure S1**). Drawing errors in the tactile experiments were similar to those in our phosphene drawing experiments (**Figure S2**).

### 2.2 Phosphene orientation is aligned with retinal nerve fiber bundles

As described above, computational models and electrophysiological evidence from *in vitro* preparations of rat and rabbit retina suggest that retinal implants may stimulate passing axon fibers (*18, 38, 39*). Retinal ganglion cells send their axons on highly stereotyped pathways *en route* to the optic nerve (**Figure 4A–C**). Because of this topographic organization, an electrode that stimulates nearby axonal fibers would be expected to antidromically activate cell bodies located peripheral to the point of stimulation. Perceptually, activating an axon fiber that passes under a stimulating electrode is indistinguishable from the percept that would be elicited by activating the corresponding ganglion cell *body*. The visual percept should appear in the spatial location in visual space for which the corresponding ganglion cell encodes information, which could be hundreds of microns away from the stimulation site (*40*). Consequently, percepts elicited from axonal stimulation would be expected to appear elongated in the direction of the underlying nerve bundle trajectory (**Figure 4C**).

**Figure 4:**
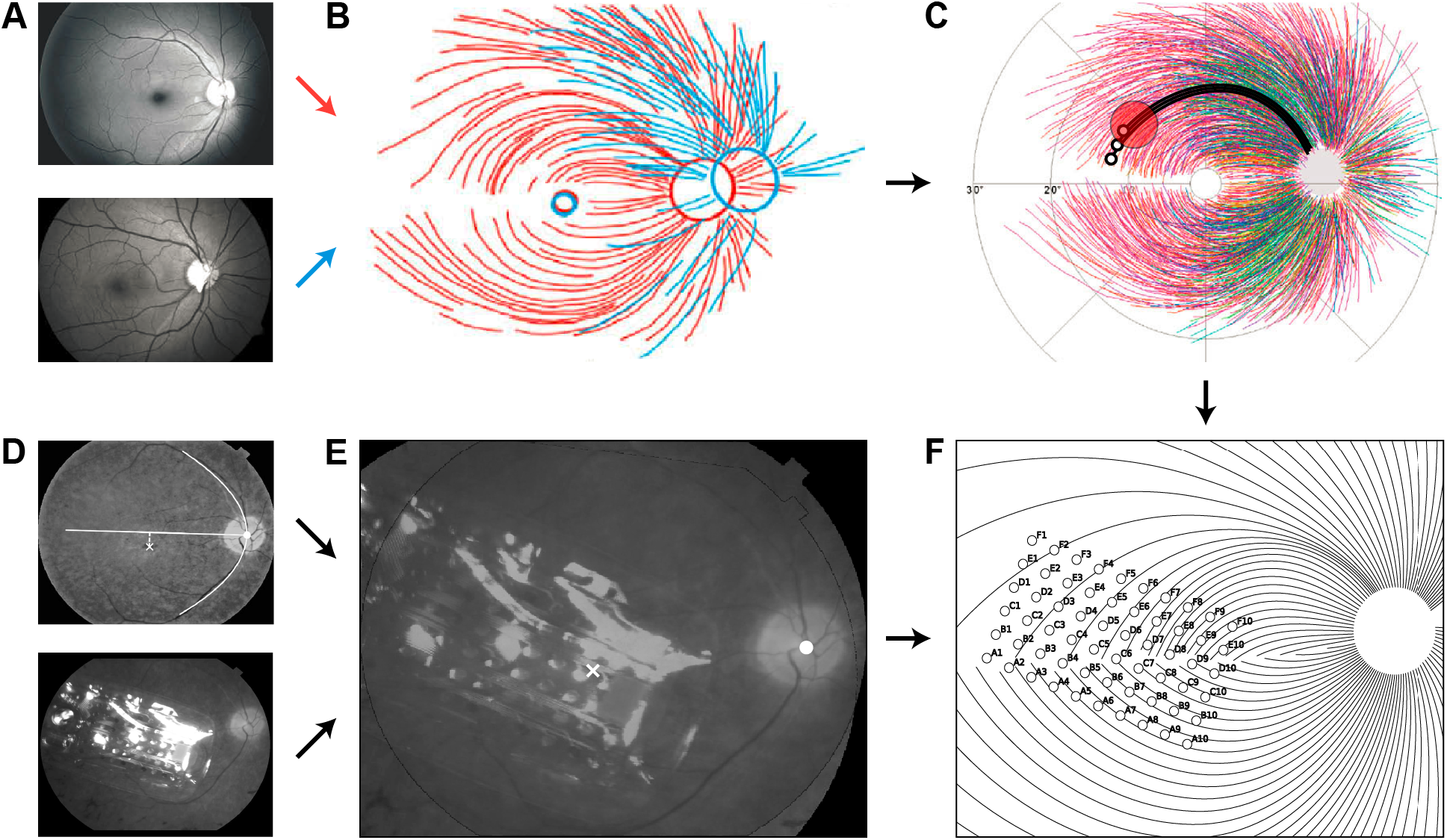
(**A–C**) The topographic organization of optic nerve fiber bundles is highly stereotyped in the human retina (adapted from 1). Fundus images from 55 human eyes (**A**) were superimposed by translation in order to center the foveola (**B**), followed by rotation and zooming to align the centers of the optic disc (**C**). Electrical stimulation (red circle) of a nerve fiber bundle could antidromically activate ganglion cell bodies peripheral to the point of stimulation (small black circles), leading to percepts that appear elongated in the direction of the underlying nerve bundle trajectory. (**D–E**) The location and orientation of each subject’s implant (Subject 4 shown here) was estimated by combining their postsurgical fundus photograph (**D**, bottom) with a baseline presurgical image in which the fovea had been identified (**D**, top) to produce a registered image (**E**; x: foveal pit, o: optic disc). The horizontal raphe (**D**, white line) was approximated by fitting a parabola to the main vascular arcade and finding the tangent to the parabola inflexion point. (**F**) The extracted landmarks were then used to place a simulated array on a simulated map of nerve fiber bundles.

To test whether the orientation of phosphene drawings were aligned with the underlying nerve fiber bundles, we estimated the relative location and orientation of each subject’s implant with respect to the fovea and the optic disc, using ophthalmic fundus photographs (**Figure 4D**; here shown for Subject 4). While yellow screening pigments allow for visualization of macular extent in normal eyes, it is difficult to discriminate the macula-periphery boundary in our subjects because of the characteristic pigmentary deposits associated with retinitis pigmentosa (*2*). We therefore had a retina specialist mark the fovea on a fundus image obtained before surgery (**Figure 4D**, top), and subsequently used computer vision techniques (see Methods) to align the presurgery image with a second fundus image obtained after surgery (**Figure 4D**, bottom), showing the implant relative to the optic nerve head. This allowed us to estimate the array center with respect to the fovea, the array rotation with respect to the horizontal raphe, and the retinal distance between the fovea and the optic nerve head for each subject (**Figure 4E**). The resulting topographic measurements were then used to simulate a map of the ganglion axon pathways (*1*) that was tailored to each subject’s retinal dimensions (**Figure 4F**).

Remarkably, we found that for all subjects, phosphene orientation was well aligned with the tangent line of the nerve fiber bundle directly below the stimulating electrode (**Figure 5**). Here, insets show mean drawings for representative electrodes, from which phosphene orientation was calculated. This was not only true for line phosphenes, which resembled carbon copies of the underlying fiber bundle topography (e.g., Subject 1: D2, Subject 4: F2), but also for more compact phosphenes, which still tended to be elongated in the direction of the local fiber bundle trajectory (e.g., Subject 2: B9, Subject 4: C10).

**Figure 5:**
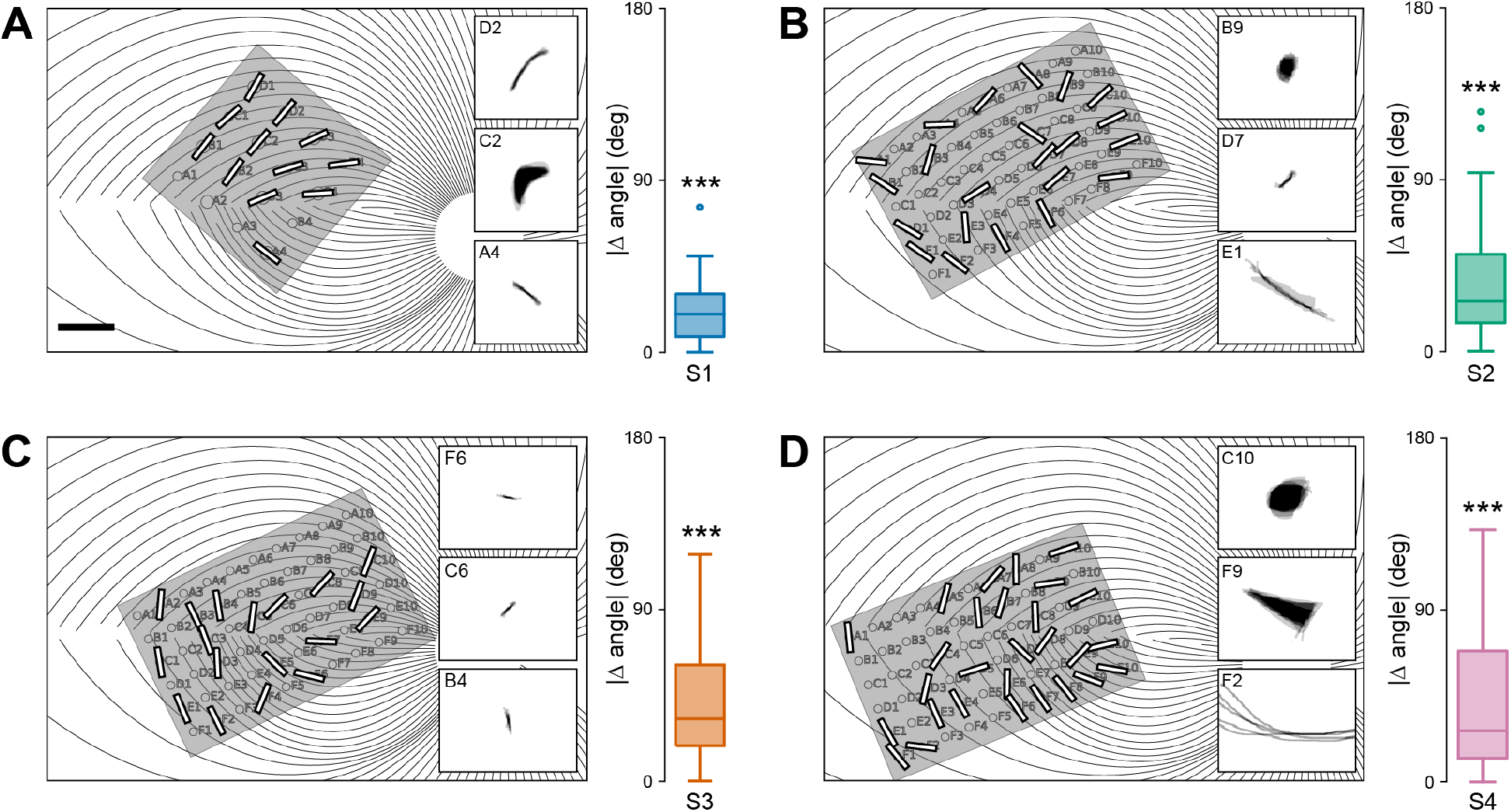
Phosphene orientations are aligned with retinal nerve fiber bundles. (**A–D**) Simulated map of nerve fiber bundles for Subjects 1–4 (scale bar: 1 mm). Phosphene orientation is indicated as oriented bars, overlaid over the corresponding electrode in the array. Insets show example percepts. Note that the maps are flipped upside down so that the upper image half corresponds to the upper visual field (inferior retina). Box plots indicate the distribution of mean absolute angular errors between phosphene orientation and the tangent line of the ganglion axon pathway nearest to the corresponding electrode. For all subjects, angular errors were significantly better than would be expected from random array placement.

To assess whether these angular errors were smaller than would be predicted from a random placement of the array on the retina, we performed a resampling analysis. First, we calculated the mean absolute angle between the five drawings corresponding to a single electrode and the tangent line of the closest nerve fiber bundle. We repeated this procedure for all electrodes in the array to produce a distribution of mean angular errors (box plots in **Figure 5**). Our resampled distribution (1000 iterations) was generated by randomly placing the array on the retina (array center coordinates: *x* ∈ [−30,30]°,*y* ∈ [−15,15]°. Probability values were estimated by comparing the mean angular error of the real distribution to the 1-tailed confidence interval of values from the resampled null model. Angular errors were significantly smaller than predicted by the null model in all four subjects (S1: *p* < 0.001, S2: *p* < 0.001, S3: *p* < 0.001, S4: *p* < 0.001).

### 2.3 Predicting phosphene shape based on a simulated map of ganglion axon pathways

We then tested whether the spatial layout of ganglion axon pathways could account for phosphene shape as well as orientation. We assumed that the activation of an axon elicited a percept centered over the receptive field location of that axon’s cell body. The activation sensitivity of a passing axon fiber was assumed to decay exponentially with retinal distance from the stimulation site, with each subject’s data being fit with two parameters: a decay constant λ, which described activation sensitivity along the axon, and a decay constant *ρ*, which described sensitivity orthogonal to the axon (see Methods). This allowed us to generate a tissue activation map for each stimulating electrode, which we thresholded to arrive at a binary image that could serve as a prediction of a phosphene drawing (small schematic in the center column of **Figure 6**).

**Figure 6:**
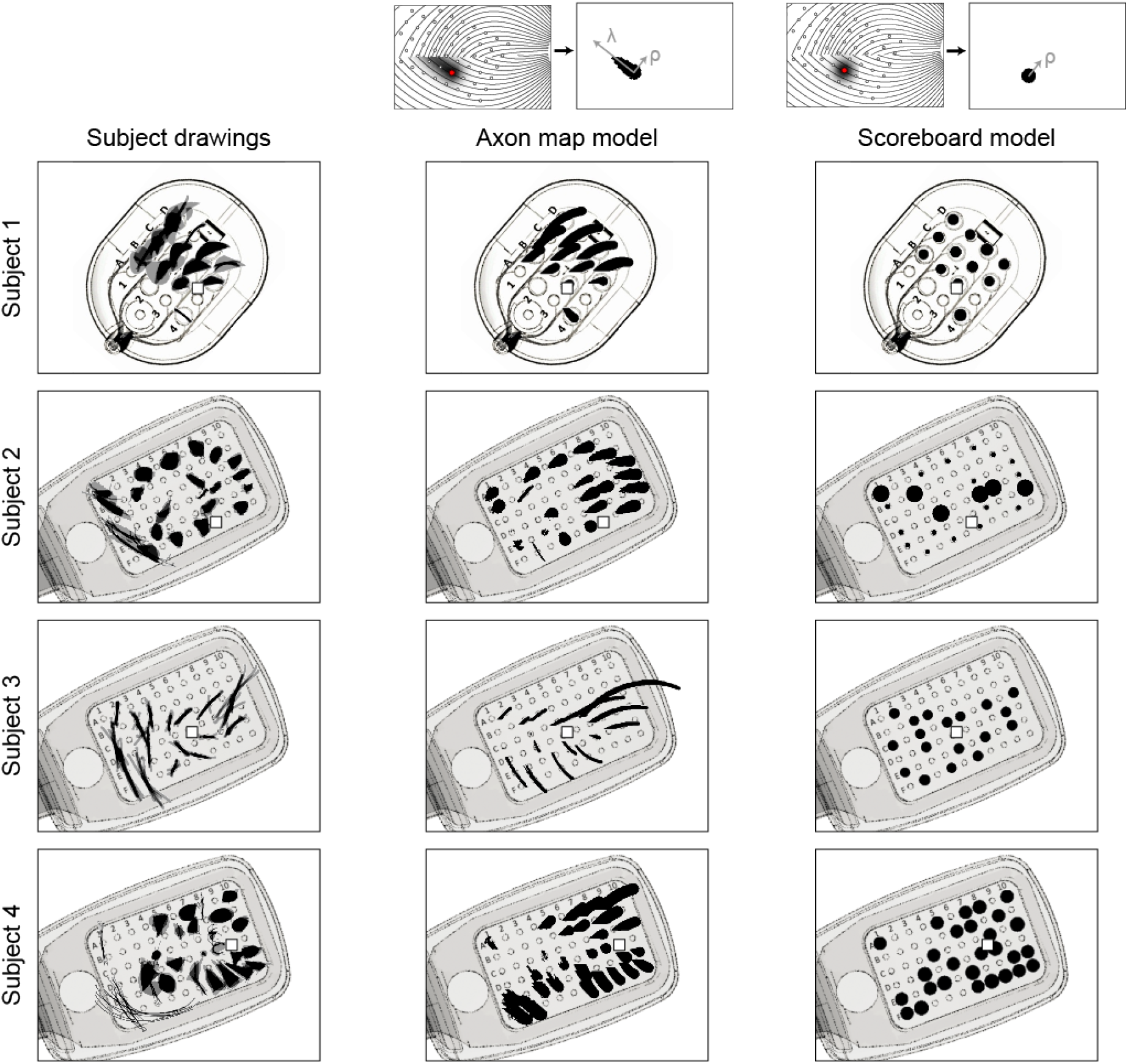
Phosphene drawings (left columns) contrasted against cross-validated phosphene predictions of the axon map model (center column) and the scoreboard model (right column), overlaid over a schematic of each subject’s implant. Each predicted phosphene is from the test fold of a leave-one-electrode-out cross-validation.

Alternatively, we considered a simpler but widely used model that treated each electrode in an array as a ‘pixel’ in an image, thus assuming that stimulating a grid of electrodes on the retina would result in the percept of a grid of isolated, focal spots of light (*33–37*). We refer to this model as the ‘scoreboard model’, because much like the large scoreboards found in sports stadiums, an image is created by an array of individual light sources that can be turned off or on. To implement the scoreboard model, we assumed that an electrode would lead to the percept of a Gaussian blob (with spatial decay constant *ρ*). The resulting intensity profile was again thresholded to obtain a binary image, which was compared to real phosphene drawings (small schematic in the right column of **Figure 6**).

To find the parameter values under each model that best predicted phosphene shape, we constructed a cost function from the difference between predicted and observed phosphene area, orientation, and elongation, which we minimized using particle swarm optimization (see Methods). Because scoreboard and axon map models had a different number of parameters (scoreboard model: *ρ*; axon map model: *ρ*, *λ*), we used leave-one-electrode-out cross-validation to allow for fair model comparison, where we repeatedly fit each model to the drawings from all but one electrode in the array. Fitted parameter values were then used to predict the phosphene shapes of the held-out drawings. Note that a single value of *ρ* and *λ* was used to describe the drawings of all electrodes in that subject’s array.

The result of this cross-validation procedure is shown in **Figure 6**. Here, ground-truth drawings are shown in the left column, and predicted phosphenes (on the test-fold of the cross-validation procedure) are shown in center and right columns. Thus, predicted phosphene shapes represent each model’s ability to generalize to new electrodes. Whereas the axon map model was able to generate both compact and elongated phosphenes that span a range of geometrical shapes such as ‘blobs’, ‘lines’, and ‘wedges’, the scoreboard model exclusively predicted round phosphenes of various size. The best fitting, cross-validated parameter values are given in **Table 1** (averaged across folds). Even though phosphene shape often varied drastically across electrodes, the axon map model recovered similar values for *ρ* and *λ* across different folds for a given subject, as indicated by relatively small SEMs. Without adjusting for drawing bias, these results suggest that electrical stimulation influences ganglion cells whose cell bodies are at a distance of approximately *ρ*=437 μm (corresponding to ~1.5° of visual angle) orthogonal to the direction of the axon fiber bundle, but as far as *λ*=1,420 μm (corresponding to ~5° of visual angle) along a direction parallel to the axon fiber.

**Table 1:** Cross-validated model parameter values, averaged across folds ± uncorrected SE.

| Subject ID | Axon map model |  | Scoreboard model |
| --- | --- | --- | --- |
| | $\rho$ ($\mu\text{m}$) | $\lambda$ ($\mu\text{m}$) | $\rho$ ($\mu\text{m}$) |
| 1 | 410 ± 5 | 1190 ± 157 | 533 ± 11 |
| 2 | 315 ± 17 | 500 ± 142 | 244 ± 34 |
| 3 | 86 ± 3 | 992 ± 150 | 170 ± 1 |
| 4 | 437 ± 6 | 1420 ± 42 | 175 ± 1 |

**Table 1a:**
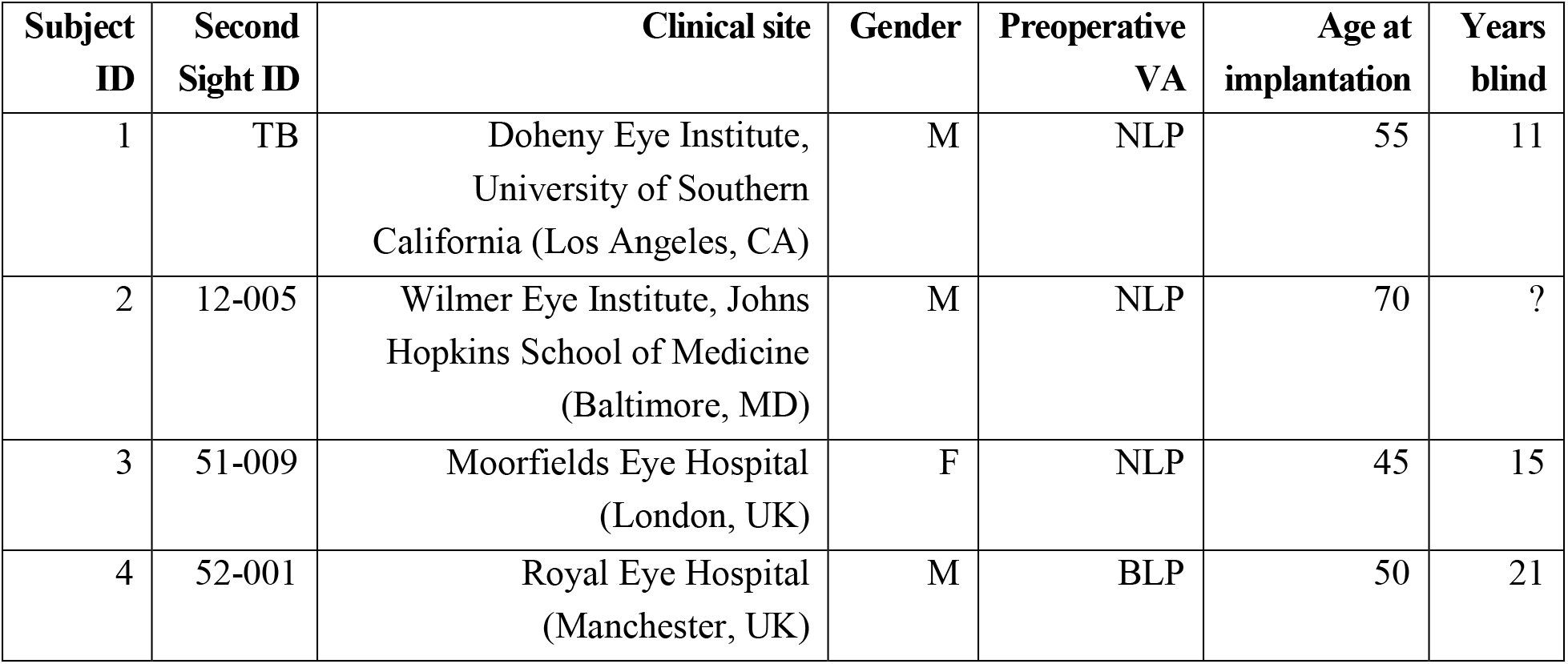
Subject details. Columns 3–7 indicate the implant site, gender, preoperative visual acuity (VA) categorized as either bare light perception (BLP) or no light perception (NLP), the age at implantation, and the number of years participants had been blind prior to implantation (self-reported). Years blind for Subject 2 is unknown due to gradual loss of vision.

To further quantify model performance, we compared cross-validated prediction errors (Equation 13) across axon map and scoreboard models, **Figure 7**. Here, each data point in the scatter plots corresponds to the cross-validated prediction error (averaged across every drawing in that fold) for each electrode. Data point almost always lie below the diagonal, indicating that the axon map model was more accurate than the scoreboard model. Indeed, the axon map model often improved cross-validated log prediction error by an order of magnitude (see insets), simply by adding a single parameter *λ* that accounted for the current spread along axons of passage in the optic nerve fiber layer of the retina.

**Figure 7:**
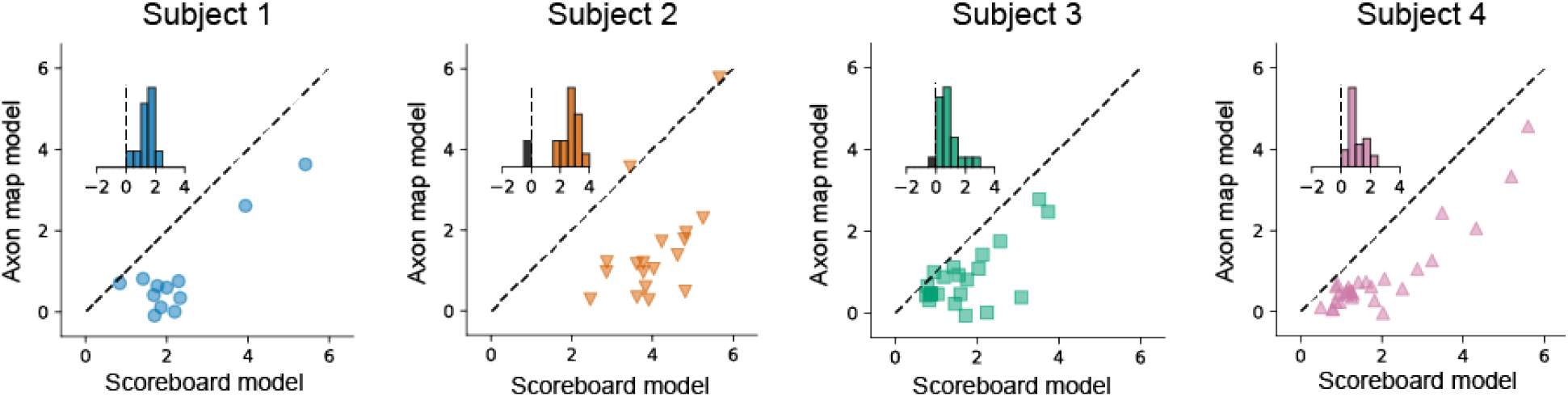
Comparison of log mean prediction error for the two models. Prediction error was based on the sum of differences between predicted and observed phosphene area, orientation, and elongation (see Equation 13). Each data point in the scatter plots corresponds to the cross-validated prediction error of all drawings associated with a particular held-out electrode. Prediction error was significantly higher for the scoreboard model compared to the axon map model (Subject 1: *p*<0.001, N=12; Subject 2: *p*<0.001, N=22; Subject 3: *p*<0.001, N=18; Subject 4: *p*<0.001, N=28; 2-tailed Wilcoxon signed-rank test). Insets in each panel show the histogram of pair-wise differences in log prediction error.

## 3. Discussion

We show here that the elicited percepts of patients with retinal implants can be accurately predicted using the spatial layout of ganglion axon pathways in the human retina. Model fits to behavioral data suggest that sensitivity to electrical stimulation is not confined to the axon initial segment (*38*), but can be modeled as falling off with different decay constants along the axon (with *λ* ranging from 500–1,420 μm) and orthogonally from the axon (with *ρ* ranging from 86–437 μm), resulting in visual percepts ranging from ‘blobs’ to ‘streaks’ and ‘wedges’ depending on both the relative values of *λ* and *ρ*, and the retinal location of the stimulating electrode. These results are in agreement with theoretical work suggesting an anisotropic spread of current in the retinal tissue (*29*) as well as previous animal literature demonstrating that epiretinal stimulation leads to activation of passing axon fibers (*18, 38, 39, 41*), which can severely distort the quality of the generated visual experience (*16, 18, 40, 42, 43*). Our findings suggest that the spatial distortions reported by patients are not arbitrary, but rather depend on the topographic organization of optic nerve fiber bundles in each subject’s retina, which can be captured by a computational model. Having an accurate model that generalizes across patients is crucial for retinal prostheses to be able to generate more complex, perceptually intelligible percepts. Our results therefore open up the possibility for future devices to incorporate stimulation strategies that are tailored to the predicted perceptual experience of each individual patient, relying on the known surgical placement of the device and empirical estimates of *ρ* and *λ*.

### 3.1 A rich repertoire of phosphene shapes

The phosphenes elicited by single-electrode stimulation vary dramatically across subjects and electrodes (**Figure 2**, **Figure 3A–C**), despite relatively small drawing errors and consistency in drawings within a given electrode (**Figure 3D–F**). These results are in agreement with the previous literature that has reported that patients subjectively report a variety of percept shapes (*23, 44–46*), of which only a small fraction could be described as focal spots of light.

The variability in phosphene shape across subjects that we report (captured by variation in *λ* and *ρ* across patients), might be due to a number of factors, a few of which are outlined below. First, diseases such as retinitis pigmentosa and macular degeneration are characterized by a progressive degeneration of photoreceptors, gradually affecting other layers of the retina (*5,47–49*). In severe end-stage retinitis pigmentosa, roughly 95% of photoreceptors, 20% of bipolar cells, and 70% of ganglion cells degenerate (*50*), so that little or no useful vision is retained. Disease progression therefore influences the relative proportion of surviving bipolar and ganglion cell types, which in turn is likely to influence phosphene shape.

Second, with a diameter of 200 μm, each electrode in the Argus II array encompasses the equivalent area of hundreds of photoreceptors. A single electrode therefore inevitably leads to activation of a wide variety of morphologically and functionally distinct retinal cells (*51, 52*), including simultaneous activation of both ON and OFF pathways. This is in contrast to natural stimulation, which precisely activates a number of specialized, functionally complementary, parallel processing pathways in the retina (for a recent review see 53). Although epiretinal stimulation with relatively short pulses might primarily activate ganglion cells rather than bipolar cells (*54–58*), there is still much to be learned about how the information from these different retinal representations are combined at later stages of processing to form a conscious percept.

Third, the mapping of retinal eccentricity to visual field coordinates is nonlinear. Because the foveola contains only photoreceptors, ganglion cell bodies are displaced centrifugally from their cone inputs by several degrees; an effect that extends out as far as 17° (*59, 60*).

Finally, phosphene size might be influenced by ganglion cell density and receptive field size. Whereas the receptive field size of retinal ganglion cells only gradually increases with eccentricity (*61*), ganglion cell density decreases rapidly (*59*). Furthermore, retinal degeneration in retinitis pigmentosa tends to progress from the periphery to the macula, thereby having a greater effect on ganglion cell density in the periphery (*47, 50, 62*). As a consequence, more peripheral electrodes would typically stimulate cells with only slightly larger receptive fields, but in much smaller numbers than in the fovea. These two conflicting effects may contribute to our finding of no correlation between phosphene area and retinal eccentricity (data not shown).

### 3.2 Phosphene shape is mediated by axonal stimulation

Despite the variability in phosphene shape, all subjects reported seeing elongated phosphenes on at least a subset of electrodes (Figure 3C). Although the electric field generated by a disk electrode is radially symmetric, the neural tissue induces anisotropies in the electric field, and stimulation of axon fibers produces even more striking anisotropies in patterns of neural activation within the retina (*29*). It has long been known that external stimulation of an axon induces an action potential that travels both backward to the cell body and forward to the synaptic terminals (*63, 64*).

A number of studies have previously hypothesized that axonal stimulation could lead to phosphenes that are elongated in shape and poorly localized (e.g., 40). However, this idea has never been explicitly tested. In the present study we demonstrate that axonal stimulation in the retina leads to predictable distortions of shape in human patients, which can be captured by a computational model (**Figures 5–7**).

Axonal stimulation is a concern for other implant technologies as well. Although subretinal prostheses such as Alpha-IMS (*9*) have electrodes in close proximity to bipolar cells, *in vitro* animal studies have found that subretinal stimulation with 1 ms pulses also directly activates retinal ganglion cells at thresholds statistically similar to those of inner retinal cells (*65–67*). Similarly, axonal stimulation is expected to be an issue for cortical implants, since passing axons from neurons located in distant parts of the brain have been shown to be highly sensitive to electrical stimulation (*68–70*).

Several recent studies have tried to identify stimulation protocols that minimize axonal activation, with mixed results. Whereas one *in vitro* study suggested using short-duration pulses (≤ 100 μs) to avoid axonal stimulation (*71*), another study did not see any benefits of short pulses, and instead suggested using long-duration pulses (≥ 25 ms) or low-frequency (< 25 Hz) sinusoidal stimulation (*39*). One difficulty with these approaches is that they are likely to limit stimulation to a highly restricted amplitude and/or frequency range, potentially limiting the dynamic range available for the encoding of brightness (*45, 46*).

We show here that percepts are highly consistent over time and can potentially be described using an anatomically detailed computational model with a small number of parameters. Thus, an alternative strategy might be to move away from thinking about artificial sight as a linear combination of ‘pixels’, and instead accept the perceptual distortions resulting from axonal stimulation as the fundamental building blocks of prosthetic vision.

## 4. Methods

### 4.1 Study design

This study was designed to investigate perceptual distortions in retinal prosthesis patients. Our goal was to systematically characterize the range of phosphene shape encountered by users of the Argus Epiretinal Prosthesis System (Second Sight Medical Products, Inc.), and to develop a computational model that could account for the reported percepts. We had access to four subjects suffering from severe retinitis pigmentosa (see Subjects) who were chronically implanted with the Argus system (see Implant Specification). Upon electrical stimulation of the retina, we asked subjects to outline perceived phosphene shape either on a grid screen or a computer touchscreen (see Psychophysical Methods). Based on ophthalmic fundus photographs, we then developed a computational model to simulate the topographic organization of optic nerve fiber bundles in each subjects’ retina (see Computational Methods). This model was used to generate predictions about the apparent shape of the expected visual percepts, which we compared to the real phosphene drawings. We concluded the study by evaluating model performance using a cross-validation procedure.

### 4.2 Subjects

Participants were four blind subjects (one female and three male) with severe retinitis pigmentosa, ranging from 45 to 70 years in age (**Table 2**). Subjects were chronically implanted with an epiretinal prosthesis as part of an FDA approved clinical trial (clinicaltrials.gov identifier for Subject 1: NCT00279500, completed; Subjects 2–4: NCT00407602, active). Surgeries were performed at the Doheny Eye Institute at the University of Southern California (Los Angeles, CA; Subject 1), at the Wilmer Eye Institute at Johns Hopkins School of Medicine (Baltimore, MD; Subject 2), at the Moorfields Eye Hospital (London, UK; Subject 3), and at the Royal Eye Hospital (Manchester, UK; Subject 4). None of the subjects had a recordable visual acuity prior to surgery, scoring worse than 2.9 logMAR (worse than 20/15,887) on a four-alternative forced-choice square-wave grating test (*19, 72*).

**Table 2:** Estimated locations of the implant and optic disc with respect to the fovea located at (0, 0) using fundus photography. Array rotation was measured with respect to the horizontal raphe.

| Subject ID | Array center (x, y; $\mu\text{m}$) | Array rotation (deg) | Optic disc center (x, y; deg) |
| --- | --- | --- | --- |
| 1 | (-651, -707) | -49.3 | (14.0, 2.40) |
| 2 | (-1331, -850) | -28.4 | (16.2, 1.38) |
| 3 | (-467, 206) | -25.8 | (14.0, 1.24) |
| 4 | (-1807, 401) | -22.1 | (16.3, 2.37) |

Due to their geographic location, Subjects 2 – 4 were not directly examined by the authors of this study. Instead, initial experimental procedures were sent to the clinical site, and trained field clinical engineers performed the experiments as specified. Raw collected data was then sent to the authors for subsequent analysis.

All tests were performed after obtaining informed consent under a protocol approved by the Institutional Review Board (IRB) at each subject’s location and under the principles of the Declaration of Helsinki.

### 4.3 Implant specification

Subject 1 was implanted with a 16-channel microelectrode array (Argus I; Second Sight Medical Products, Inc., Sylmar, CA) consisting of 260 and 520 μm diameter platinum disc electrodes, subtending 0.9° and 1.8° of visual angle, respectively. Electrodes were spaced 800 μm apart, and arranged in a 4×4 alternating checkerboard pattern (**Figure 1A**). Subjects 2 – 4 were implanted with a 60-channel microelectrode array (Argus II; Second Sight Medical Products, Inc., Sylmar, CA) consisting of 200 μm diameter platinum disc electrodes, each subtending 0.7° of visual angle. Electrodes were spaced 525 μm apart, and arranged in a 6×10 grid (**Figure 1B**).

All stimuli described in this study were presented in ‘direct stimulation’ mode. Stimuli were programmed in Matlab using custom software, and pulse train parameters (the electrode(s) to be stimulated, current amplitude, pulse width, individual pulse duration, inter-pulse interval, and pulse train duration) were sent directly to an external visual processing unit (VPU), which was used to send stimulus commands to the internal portion of the implant using an inductive coil link (‘camera’ mode). The implanted receiver wirelessly received these data and sent the signals to the electrode array via a small cable.

In day-to-day use, an external unit consisting of a small camera and transmitter mounted on a pair of glasses is worn by the user. The camera captured video and sends the information to the VPU which converts it into pulse trains using pre-specified image processing techniques.

### 4.4 Psychophysical methods

Perceptual thresholds for individual electrodes were measured using an adaptive yes/no procedure implemented using custom software (see Supplemental Information). All presented stimuli were charge-balanced, square-wave, biphasic, cathodic-first pulse trains with fixed stimulus duration (Argus I: 500 ms, Argus II: 250 ms), current amplitude (set at 2x threshold), stimulus frequency (20 Hz) and pulse duration (0.45 ms/phase, no interphase delay).

Subjects were asked to perform a drawing task with a tactile target (Supplemental Information) or when their retina was electrically stimulated (**Figure 1**). In a given experimental run, a total of *n* stimulus conditions (either tactile or retinal stimulation) were tested. Each condition was repeated for *m* trials (for a total of *mn* trials per experimental run). Repeated trials of the same condition were randomized amongst other stimuli to confirm reproducibility of results.

Head movement of the Argus I subject was minimized with a chin rest. After each stimulus presentation, the subject traced the shape on a grid screen (containing 6 inch horizontal and vertical grid lines) with a center location aligned horizontally and vertically with the subject’s head. Drawing was carried out with a pen whose cap was a different color than its body. A head-mounted camera (Misumi CMOS S588-3T), located on the subject’s glasses, was used to record the trials to digital video recorder (DVR). Video files were analyzed off-line to extract shape data using custom built tracking software. In the first stage of processing, the entire image was rotated appropriately using the grid screen background as a reference. In the second stage, vertical and horizontal gridlines, and the distance from the subject to screen were used to set a new coordinate system in visual angle coordinates (since the subject was 16 inches / 40.6 cm from the screen, 4 gridlines = 70.0 cm corresponded to 73.8 degrees visual angle). In the third stage, the location of the pen cap was tracked (based on its color) across each frame of the video file. Finally, a binary shape data file was built from pen cap coordinate locations across all frames.

Argus II subjects were placed in a chair at a comfortable distance from a touch screen monitor with its center location aligned horizontally with the subject’s head. The distance from each subject’s eyes to the screen was recorded. After each stimulus presentation, the subject traced the shape on the monitor and the experimenter advanced to the next trial. Touch screen data were instantly recorded by custom software in 2D coordinates to a text file. Text files were analyzed offline to translate vector coordinates to a binary shape data file. The distance recorded from the subject to screen was used to set a new coordinate system in visual angle. Since Subjects 2 – 4 were 33, 30.0, and 30.5 inches from the screen, this corresponded to a display size of 60, 65 and 64 degrees of visual angle (horizontal screen length), respectively. After translating to the final visual angle coordinate system, the binary image was used in subsequent shape analyses.

### 4.5 Computational methods

#### 4.5.1 Phosphene shape descriptors

Phosphene shape was quantified using three parameter-free shape descriptors commonly used in image processing: area, orientation, and elongation (*73*). (Elongation is sometimes also referred to as eccentricity in the literature. We avoid that usage here to prevent confusion with retinal eccentricity). These descriptors are based on a set of statistical quantities known as ‘image moments’. For an *X* × *Y* pixel grayscale image, *I*(*x,y*), where *x* ∈ [1,*X*] and *y* ∈ [1,*Y*], the raw image moments *M_ij_* were calculated as:

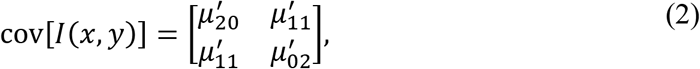

Raw image moments were used to compute area (*A* = *M*_00_) and the center of mass 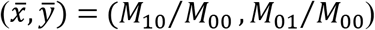 of each phosphene.

Phosphene orientation was calculated from the covariance matrix of an image:

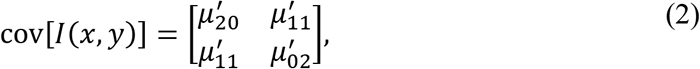

where 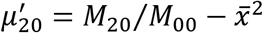, 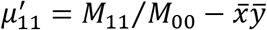 and 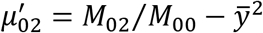. The eigenvectors of this matrix corresponded to the major and minor axes of the image intensity. Orientation (*θ*) could thus be extracted from the angle of the eigenvector associated with the largest eigenvalue towards the axis closest to this eigenvector:

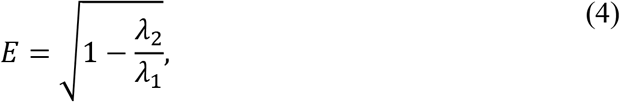

which was valid as long as 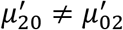, with *θ* ∈ {– *π*/2, *π*/2}. To avoid division by zero, we manually assigned an angle of *θ* = 0 whenever 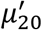 was equal to 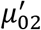.

Phosphene elongation (*E*) was calculated from the eigenvectors of the covariance matrix of Equation (2):

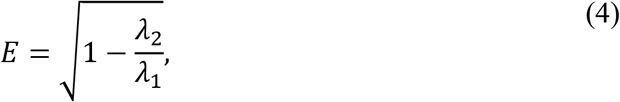

where 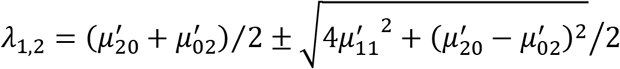, and *E* ∈ [0,1]. An elongation of *E* = 1 represents a circle, and *E* = 0 represents an infinitesimally thin line.

#### 4.5.2 Determination of implant location using fundus photography

Implant location was estimated by analyzing stereoscopic color fundus photographs obtained using systems available at each clinical site. For each subject, we performed the following procedure:

1. Extract landmarks: On a baseline fundus photograph (before surgery), a retina specialist marked the foveal pit and the center of the optic nerve head. On the most recent fundus photograph (after surgery), we marked the center of the implant.
2. Combine baseline image with implant image: We performed image registration using feature matching to bring the two images into the same coordinate system.
3. Adjust for magnification: Pixel distances were converted to retinal distances by using the known electrode-electrode spacing (Argus I: 800 μm, Argus II: 525 μm).
4. Adjust for rotation: We approximated the horizontal raphe by fitting a parabola to the main vascular arcade, assuming that the center of the optic nerve head lied at the vertex of the parabola, and that the raphe was parallel to the parabola’s axis of symmetry (*74, 75*).
5. Coordinate transform: The registered image was rotated so that the horizontal raphe came to lie on the abscissa, and the foveal pit at the origin of the new coordinate system. We located the coordinates of the center of the optic nerve head as well as the center of the array (from Step 1) in this new coordinate system. Retinal distances (μm) were related to eccentricity (deg) using a formula that computes the relationship between retinal arc lengths and visual angles from based on the optic axis (*60*).
6. The extracted landmarks were then used to place a simulated array on a simulated map of ganglion axon pathways using the pulse2percept software (*76*).

This procedure allowed us to estimate each subject’s array location and orientation with respect to the fovea (**Table 3**). Based on fundus photographs of 104 sighted humans (*77*), the center of the optic disc was expected to be located at 15.5° ± 1.1° nasal, 1.5° ± 0.9° superior with respect to the fovea. For all four subjects, the estimated center of the optic disk was within two standard deviations of these expected values.

#### 4.5.3 Scoreboard model

The scoreboard model assumed that electrical stimulation led to the percept of focal dots of light, centered over the visual field location associated with the stimulated retinal field location (*x*_stim_, *y*_stim_), whose spatial intensity profile decayed with a Gaussian profile (*36, 37*):

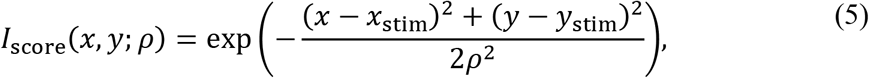

where *ρ* was the spatial decay constant.

The resulting intensity profile *I*_score_(*x*,*y*; *ρ*) was then thresholded to obtain a binary image. The threshold was chosen as 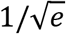, such that points closer than *ρ* to (*x*_stim_, *y*_stim_) were assigned a value of 1, and all other points were assigned a value of 0.

#### 4.5.4 Axon map model

Following Jansonius et al. (1), we assumed that the trajectories of the optic nerve fibers could be described in a modified polar coordinate system (*r, ϕ*) with its origin located in the center of the optic disc. A nerve fiber was modeled as a spiral:

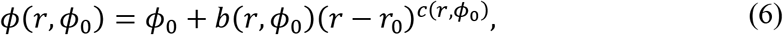

where *ϕ*_0_ = *ϕ*(*r* = *r*_0_) is the angular position of the trajectory at its starting point at a circle with radius *r*_0_ around the center of the optic disc, *b* a real number describing the curvature of the spiral,

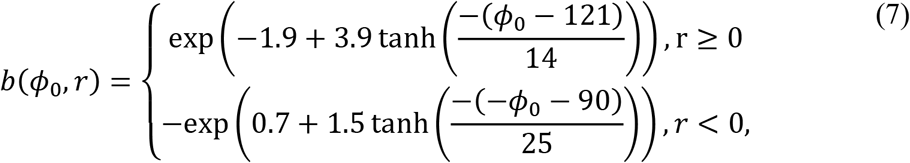

and *c* a positive real number describing the location of the point of maximal curvature,

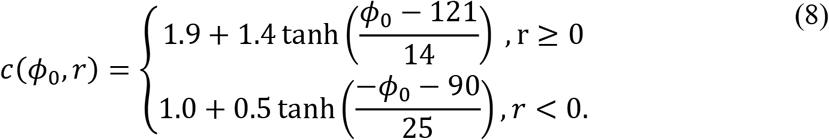

Jansonius and colleagues determined parameter values by fitting Equations (6–8) to the topographical layout of 55 eyes from 55 human subjects (for details see *1*). The attentive reader might notice that Equation (8) above fixes a typo in Equation (3) of Jansonius et al. (*1*): The tanh numerator should indeed read *ϕ*_0_ – 121, not – (*ϕ*_0_ – 121).

To apply the axon map to the eyes of our subjects, we first transformed the original coordinate system (*r, ϕ*)to Cartesian coordinates (*x, y*) with the foveal pit located at (0, 0), and then set the coordinates of the optic disc (*x*_od_, *y*_od_) to the values estimated from fundus photography (**Table 3**). The resulting axon maps for each subject can be seen in **Figure 4**.

An axon’s sensitivity to electrical stimulation was assumed to decay exponentially with distance from the soma (*x*_soma_, *y*_soma_):

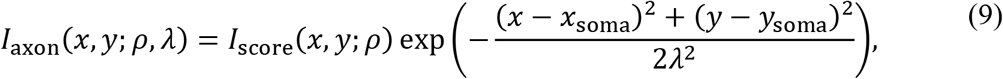

where *λ* was the spatial decay constant along the axon. *I*_score_(*x, y*; *ρ* is the same as in Equation (5) and is parameterized by a single parameter, *ρ*. As in the scoreboard model, the resulting intensity profile *I*_axon_(*x,y*; *ρ, λ*)was thresholded to obtain a binary image.

#### 4.5.5 Model fitting and evaluation

To fit the scoreboard and axon map models to subject drawings, we first calculated the coefficient of determination (*R*^2^)from the predicted binary images and the corresponding ground-truth subject drawings. *R*^2^ was calculated from the ratio of the residual sum of squares (*SS*_res_) and the total sum of squares (*SS*_tot_) for each shape descriptor (area, orientation, or elongation):

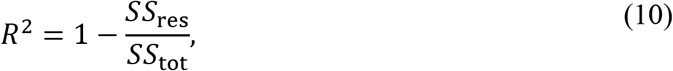

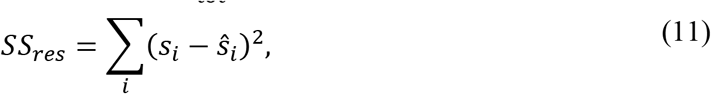

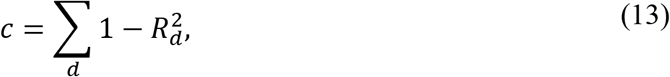

where *S_i_* was the shape descriptor for the *i*-th ground-truth image, *ŝ_i_* was the shape descriptor for the *i*-th predicted image, and š was the mean of the shape descriptor averaged over all images.that a single value The three quantities 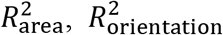, and 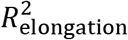 resulting from this procedure were then combined to construct a cost function that could be iteratively minimized:

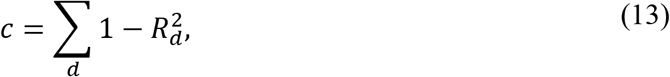

where d = {area, orientation, elongation}. Due to the nonconvexity of this optimization problem, we minimized the cost function using particle swarm optimization (*78*). We set the swarm size at ten times the number of parameters (*79*). We ran every fitting procedure five times with different, randomly chosen initial conditions, and then chose the best run in subsequent analyses.

To allow for a fair performance comparison despite the scoreboard and axon map models having different numbers of parameters, we implemented a leave-one-electrode-out cross-validation procedure, where we repeatedly fit each model to the drawings from all but one electrode in the array. This is equivalent to calculating the Akaike Information Criterion that takes into account the difference in number of parameters (*80*). The fitted parameter values were then used to predict the shape descriptors of the held-out drawings (**Figure 13**). Note that a single value of *ρ* and *λ* was fitted for each subjects, and then used for all electrodes in that subject’s array.

## 7. Acknowledgments

The authors would like to thank retina specialists Drs. Aaron Y. Lee and Cecilia S. Lee in the Department of Ophthalmology at the University of Washington for locating the fovea in fundus photographs.

## Author contributions

J.D.W., A.R., G.M.B., and I.F. designed the study. D.N. collected the data. M.B. and A.R. analyzed the data. M.B. wrote the software. M.B., D.N., and I.F. wrote the manuscript.

## Data and software availability

Data are available on the Open Science Framework (doi:10.17605/osf.io/dw9nz). The software used for analyses was based on pulse2percept (76). Scripts used to fit the scoreboard and axon map models, to analyze the data, and to produce the figures in the paper are available on GitHub (https://github.com/VisCog/ArgusShapes.git, v0.1).

## 8. Supplementary Methods

### 8.1 Threshold measurement

Perceptual thresholds were used to determine the suprathreshold level of stimulation (2x threshold) used for the drawing task described in the main text.

#### 8.1.1 Argus I subject

Perceptual thresholds were measured using custom-developed software on single electrodes using an adaptive yes/no procedure. Stimuli for measuring threshold were charge-balanced, 0.45 ms / phase, cathodic-first, biphasic 20 Hz pulse trains, 500 ms in duration. There was no interphase interval (i.e., the total cathode-anode pulse width was 0.90 ms). On each trial, subjects were asked to judge whether or not they saw a phosphene on that trial. Half of the trials were stimulus-absent catch trials interleaved randomly with the stimulus-present trials. Current amplitude was varied using a three-up-one-down staircase procedure to find the threshold current amplitude needed for the subjects to see the stimulus on 50% of stimulus-present trials, corrected for the false alarm rate. During each staircase, only amplitude varied while all other parameters (frequency, pulse width, pulse train duration, and the number of pulses) was held constant. Each threshold was based on fitting a Weibull function to a minimum of 125 trials and error bars are estimated using Monte-Carlo simulation (*81*).

#### 8.1.2 Argus II subjects

Perceptual thresholds were measured using custom-developed software using a yes/no procedure that was a hybrid between an adaptive procedure and a method of constant stimuli. Stimuli were charge-balanced, 0.45 ms / phase, cathodic-first, biphasic 20 Hz pulse trains, 250 ms in duration. As in the Argus I stimulation, there was no interphase interval (i.e., the total cathode-anode pulse width was 0.90 ms). However, in the Argus II procedure, rather than testing each electrode individually, as many as six different electrodes were tested in a single experimental run. The entire experimental run was divided into five separate blocks of 12 trials per electrode in each block for a total of 72 stimulation trials per block. In addition to stimulation trials, 32 catch trials in total were interspersed randomly over the five presentation blocks. Thus, the maximum number of trials over 5 blocks was 72×5+32 = 392 trials. Stimulus amplitudes (for stimulus present trials) for the first block were predetermined (method of constant stimuli). After the first block, a maximum likelihood algorithm fit of a Weibull function to the current data determined the range of the next block of stimulation amplitude values for each electrode. After each block a confidence interval was acquired for each electrode using a Monte-Carlo simulation based on responses to the previous trials. If the confidence interval for an electrode fell below to a pre-set level, trials for that electrode were no longer presented, but trials on the other electrodes continued through a maximum of five blocks.

Results were deemed unreliable if the false alarm rate, determined by the percentage that the subject saw a stimulus during catch trials, was greater than 20%. Data from runs with higher false alarm rates than 20% were removed from analysis and the runs were repeated.

### 8.2 Tactile target control task

Our experiments relied on the ability of our subjects to draw percepts accurately and consistently across trials. However, our blind subjects lacked tactile-visual feedback for many years. As a result, we were concerned that they might be unable to reliably reproduce visual phosphenes.

We therefore carried out a control experiment with tactile targets to establish baseline drawing errors for each subject. For the Argus I subject (Subject 1), the test stimuli consisted of a set of 11 tactile shapes made of felt with a cardboard background (**Figure S1A**). For Argus II subjects, methods were refined to use six tactile shapes made of felt with a cardboard background (**Figure S1B**). Subjects were asked to feel the felt shapes (**Figure S1C**), and then draw them on a board or a touch screen.

**Figure S1.**
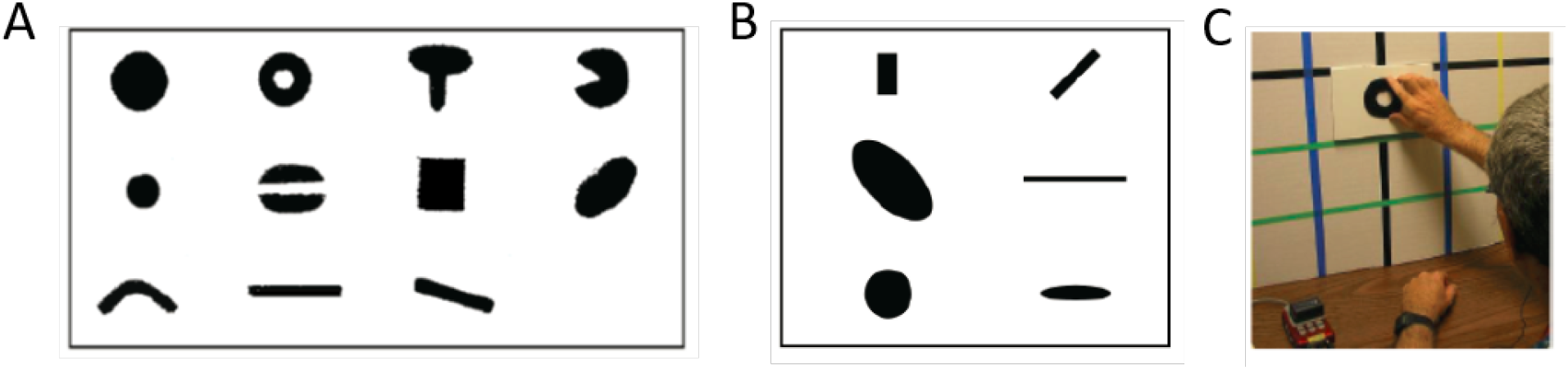
Tactile target control task. (**A**) Argus I tactile shapes. (**B**) Argus II tactile shapes. (**C**) Subject feels shape before tracing on a screen.

Shapes were classified in terms of their area, their orientation, and the major and minor axis lengths calculated from raw image moments. Visual inspection of the drawings suggested that subjects may differ in their drawing ability between compact and elongated shapes. We therefore subdivided the data into two subgroups (‘elongated’: major axis length larger than twice the minor axis length, equivalent to elongation 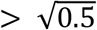 in the main text), and plotted drawing errors separately.

We calculated drawing errors as the differences across repeated drawing trials for a given tactile target (**Figure S2**). Subject 1 showed less area variability across trials for compact (17±2% error) than for elongated shapes (34±2% error). Orientation was less variable for elongated (8°±2° error) than for compact shapes (22°±4° error). Subject 2 showed similar area variability for compact (20±5% error) and elongated shapes (17±9% error). Orientation was less variable for elongated (6°±3° error) than for compact shapes (23°±20° error). Subject 3 showed comparable area variability for compact (20±8% error) and elongated shapes (23±3% error). Orientation was less variable for elongated (8°±2° error) than for compact shapes (22°±4° error). Subject 4 showed less area variability for compact (17±2% error) than for elongated shapes (34±2% error). Orientation variability was comparable for compact (9°±9° error) and elongated shapes (8°±6° error).

**Figure S2.**
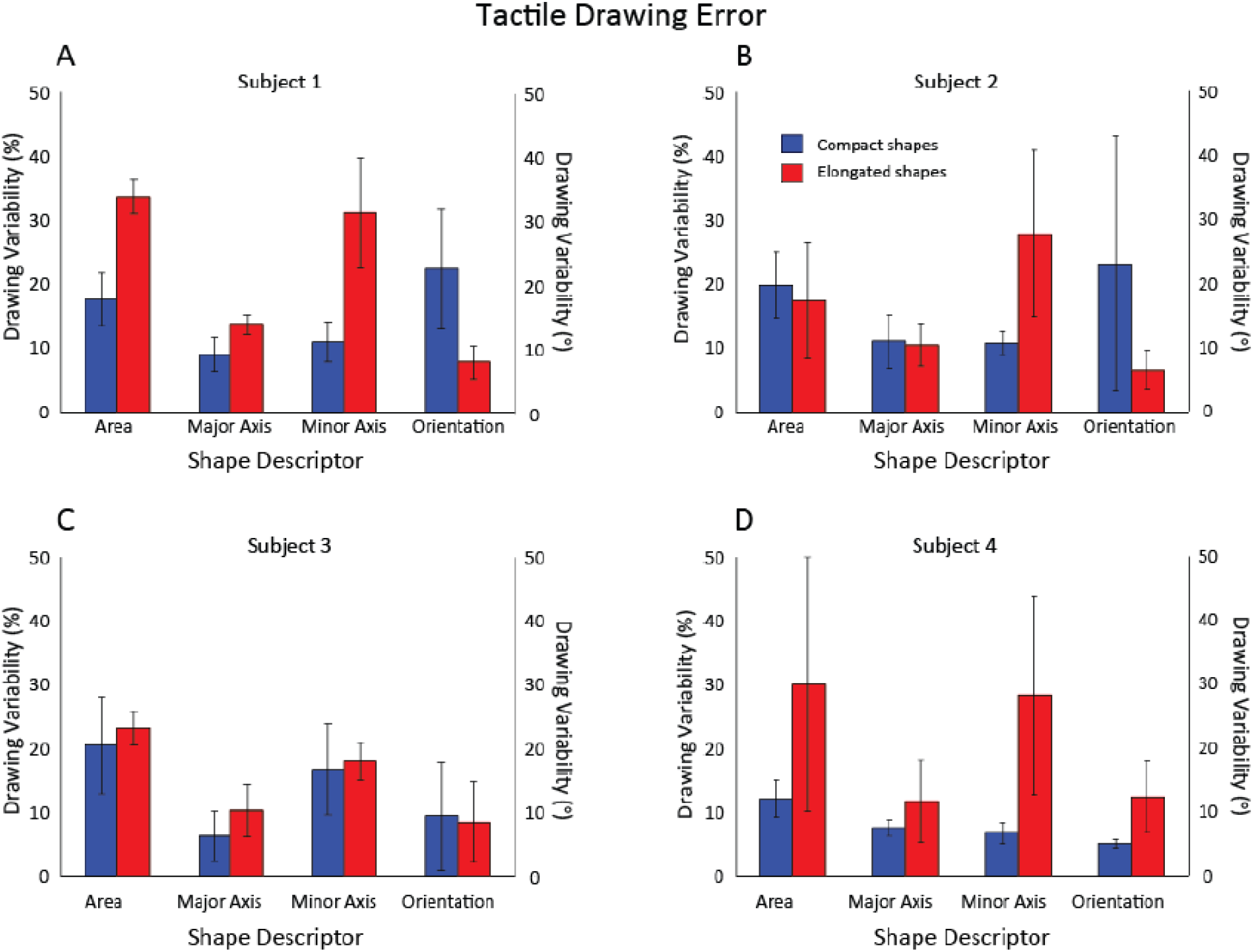
Tactile drawing errors for all subjects.

We then calculated drawing bias as the ratio or difference between the tactile target and the mean shape of drawings of that tactile target (**Figure S3**). Subject 1 drew both compact and elongated shapes larger than the actual size of the tactile targets (compact: 1.5±0.15 times larger; elongated: 1.8±0.3 times larger), and tended to draw elongated shapes biased by 9°±1.85° counter-clockwise. Subject 2 drew both compact and elongated shapes smaller than the tactile targets (compact: 0.46±0.11 times smaller; elongated: 0.56±0.09 times smaller), and tended to draw elongated shapes biased by 16°±5.41° counter-clockwise. Subject 3 drew both compact and elongated shapes larger than the tactile targets (compact: 1.5±0.31 times larger; elongated: 2.4±0.19 times larger), and tended to draw elongated shapes biased by 4°±1.7° clockwise. Subject 4 drew both compact and elongated shapes larger than the tactile targets (compact: 2.5±0.36 times larger; elongated: 2.3±0.68 times larger), and tended to draw elongated shapes biased by 14°±4.3° counter-clockwise.

Overall, trial-by-trial variability in the phosphene drawing experiments was similar to that of the tactile experiments— consistent with the data in the main paper suggesting that electrically stimulated percepts might be relatively consistent across trials.

**Figure S3.**
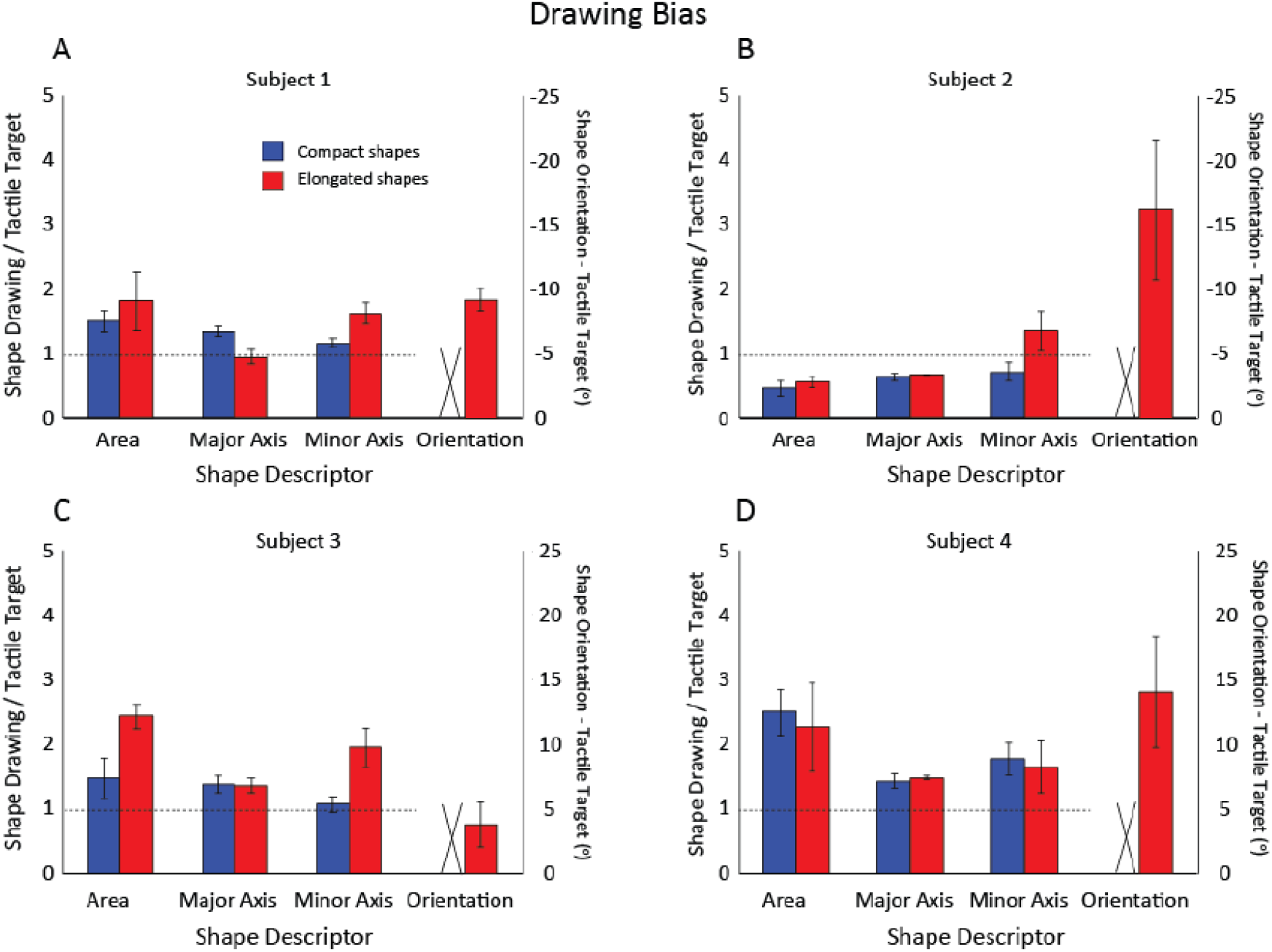
Tactile drawing biases for all subjects.

